# MARCH8 targets cytoplasmic lysine residues of various viral envelope glycoproteins

**DOI:** 10.1101/2021.04.20.440588

**Authors:** Yanzhao Zhang, Seiya Ozono, Takuya Tada, Minoru Tobiume, Masanori Kameoka, Satoshi Kishigami, Hideaki Fujita, Kenzo Tokunaga

## Abstract

The host transmembrane protein MARCH8 is a RING finger E3 ubiquitin ligase that downregulates various host transmembrane proteins, such as MHC-II. We have recently reported that MARCH8 expression in virus-producing cells impairs viral infectivity by reducing virion incorporation of not only HIV-1 envelope glycoproteins but also vesicular stomatitis virus G-glycoprotein through two different pathways. However, the MARCH8 inhibition spectrum remains largely unknown. Here, we investigate the antiviral spectrum of MARCH8 using HIV-1 pseudotyped with a variety of viral envelope glycoproteins. Pseudotyping experiments revealed that viral envelopes derived from the rhabdovirus, arenavirus, coronavirus, and togavirus (alphavirus) families were sensitive to MARCH8-mediated inhibition. Lysine mutations at the cytoplasmic tails of rabies virus-G, lymphocytic choriomeningitis virus glycoproteins, SARS-CoV and SARS-CoV-2 spike proteins, and Chikungunya virus and Ross River virus E2 proteins conferred resistance to MARCH8. Immunofluorescence showed impaired downregulation of the mutants of these viral envelopes by MARCH8, followed by lysosomal degradation, suggesting that MARCH8-mediated ubiquitination leads to intracellular degradation of these envelopes. Indeed, rabies virus-G and Chikungunya virus E2 proteins proved to be clearly ubiquitinated. We conclude that MARCH8 has inhibitory activity on a variety of viral envelope glycoproteins whose cytoplasmic lysine residues are targeted by this antiviral factor.

## Introduction

Membrane-associated RING-CH (MARCH) 8 is one of eleven MARCH family members of RING-finger E3 ubiquitin ligases and consists of a short ectodomain connecting two transmembrane domains with two cytoplasmic amino and carboxyterminal domains (Bartee et al., 2004; Goto et al., 2003). MARCH8 is known to downregulate various types of host transmembrane proteins, including MHC-II (Jahnke et al., 2013; Ohmura-Hoshino et al., 2006), CD86 (Baravalle et al., 2011; Tze et al., 2011), CD81 (Bartee et al., 2010), CD44 (Bartee et al., 2010; Eyster et al., 2011), TRAIL receptor 1 (van de Kooij et al., 2013), CD98 (Eyster et al., 2011; Funakoshi et al., 2014), Bap31 (Bartee et al., 2010), IL-1 receptor accessory protein (Chen et al., 2012), transferrin receptor (Fujita et al., 2013), and cadherin-1 (Singh et al., 2021). We have recently reported that MARCH8 downregulates HIV-1 envelope glycoproteins (Env) from the cell surface, thereby reducing the infectivity of viruses produced from MARCH8-expressing cells (Tada et al., 2015). Importantly, the antiviral effect of MARCH8 was observed in not only HIV-1 Env but also vesicular stomatitis virus G-glycoprotein (VSV-G), which was even more sensitive to this host protein (Tada et al., 2015). Furthermore, we have recently shown that MARCH8 inhibits VSV-G- or HIV-1 Env-mediated viral infectivity by two different mechanisms through ubiquitination-dependent or tyrosine motif-dependent pathways, respectively (Zhang et al., 2020). Given that these two viral envelope glycoproteins are structurally and phylogenetically unrelated to each other, we conjecture that MARCH8 might show broad-spectrum inhibition of enveloped viruses by downregulating viral envelope glycoproteins, exactly as reported for the aforementioned host membrane proteins. Here, we show that MARCH8 can target a variety of viral envelope glycoproteins by performing not only pseudotyping assays but also whole-virus-based infection experiments.

## Results

### MARCH8 expression in producer cells inhibits infection by pseudoviruses harboring various viral envelope glycoproteins

To investigate the antiviral spectrum of MARCH8, we used luciferase-reporter/HiBiT-tagged HIV-1 (Ozono et al., 2020) pseudotyped with a variety of viral envelope glycoproteins (Figure 1A). Because VSV, whose glycoprotein is highly susceptible to MARCH8-mediated inhibition (Tada et al., 2015), belongs to the rhabdovirus family, we first tested another rhabdovirus-derived envelope, rabies virus (RABV)-G. Viral infectivity assays showed that this virus was sensitive to MARCH8 in a dose-dependent manner (Figure 1B). We next tested the prototypic arenavirus lymphocytic choriomeningitis virus (LCMV) envelope GP, which carries a single transmembrane domain, along with the aforementioned viral envelopes. LCMV-GP was found to be particularly sensitive, even in the presence of only a small amount of MARCH8 expression plasmid (Figure 1C). Other viral envelopes with a single transmembrane domain, i.e., severe acute respiratory syndrome-associated coronavirus (SARS-CoV) and SARS-CoV-2 spike (S) proteins, were also dose-dependently inhibited by MARCH8 (Figure 1D). Alphavirus envelopes are initially expressed as polyproteins with multiple transmembrane domains. Chikungunya virus (CHIKV) envelope was found to be especially highly susceptible to MARCH8, whereas another alphavirus species, the Ross River virus (RRV) envelope, was also dose-dependently inhibited (Figure 1E).

**Figure 1.**
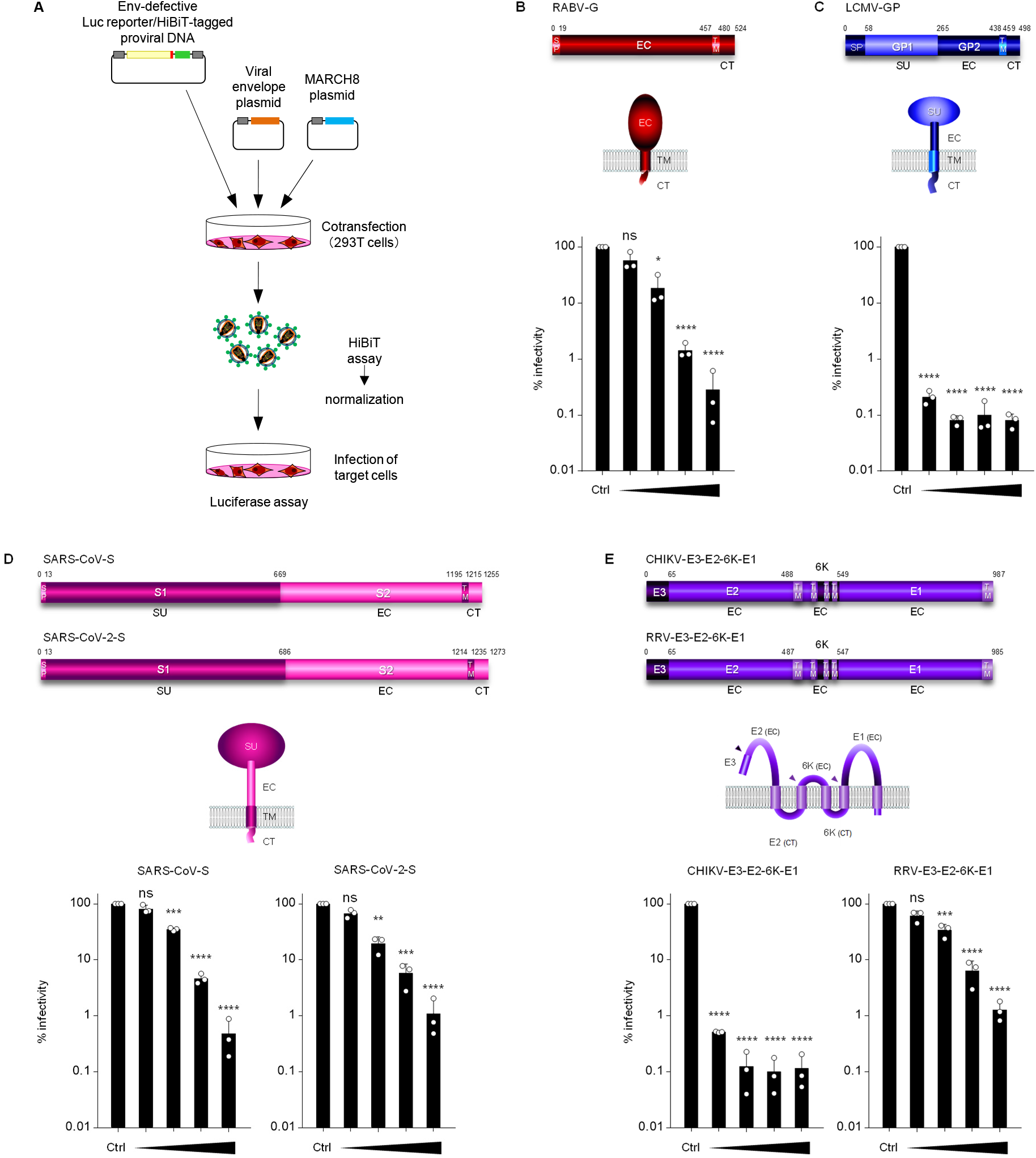
MARCH8 expression in producer cells decreases the infectivity of viruses pseudotyped with a variety of viral envelope glycoproteins. **(A)** Schematic flowchart of the experimental procedure for pseudovirus infectivity assays. Pseudoviruses were produced from 293T cells cotransfected with Env-defective HiBiT-tagged HIV-1 luciferase (luc) reporter proviral DNA clone pNL-Luc2-IN/HiBiT-E(-) and either a control (Ctrl) or increasing amounts of the MARCH8 plasmid (30, 60, 120, and 240 ng) together with various viral envelope plasmids. Produced viruses were subjected to HiBiT assays to determine the level of virion production. Equivalent amounts of p24 antigen (translated from HiBiT-luc activities) of pseudoviruses were used for the infection of different target cells, as described below. After 48 h, cells were lysed, and firefly luc activities were measured to determine viral infectivity. **(B-E)** Various viral envelope glycoproteins tested are differentially susceptible to MARCH8-mediated inhibition. Viral infectivity was determined from infection by viruses pseudotyped with **(B)** rabies virus G (RABV-G) in 293T cells; **(C)** lymphocytic choriomeningitis virus (LCMV) envelope GP in NIH3T3 cells; **(D)** severe acute respiratory syndrome-associated coronavirus (SARS-CoV) and SARS-CoV-2 spike (S) in 293T cells coexpressing ACE2 receptor and the transmembrane protease TMPRSS2; and **(E)** alphavirus E3-E2-6K-E1 (from Chikungunya virus (CHIKV) and Ross River virus (RRV)) in HeLa cells. Data from three independent experiments are shown as a percentage of the infectivity of viruses produced without MARCH8 (mean ± s.d., *n* = 3 technical replicates). \**p*<0.05, \*\**p*<0.005, \*\*\**p*<0.0005, \*\*\*\**p*<0.0001 compared with the Ctrl using sing two-tailed unpaired *t*-tests. ns, not significant. Schematic representations of genes and topological structures illustrating viral envelope glycoproteins are shown in the upper and middle panels. SP, signal peptide; EC, extracellular domain; TM, transmembrane domain; CT, cytoplasmic tail; SU, surface subunit. Arrowheads in the middle panel of E represent cleavage sites of alphavirus E3-E2-6K-E1.

### MARCH8 targets cytoplasmic lysine residues of viral envelopes with single transmembrane domains

We have recently reported that the cytoplasmic lysine residues of VSV-G are targeted by MARCH8 for ubiquitination and degradation (Tada et al., 2015), which prompted us to address whether MARCH8-sensitive viral envelopes also harbor cytoplasmic lysines that could be ubiquitination targets for MARCH8. Among the above viral envelopes, a single-spanning membrane protein, RABV-G, harbors three lysine residues at positions 489, 508, and 517 in its cytoplasmic tail. Therefore, we mutated these residues to arginines in this viral envelope (K489/508/517R) (Figure 2A). Infectivity assays to compare the WT and mutant envelopes showed that the lysine mutations in RABV-G conferred resistance to MARCH8 (Figure 2E). The cytoplasmic tail of the single-spanning membrane protein LCMV-GP contains six lysine residues at positions 465, 471, 478, 487, 492, and 496, which we mutated to arginine residues (K465/471/478/487/492/496R) (Figure 2B). The effect of the mutation in LCMV-GP was more drastic in that the mutant envelope was resistant to MARCH8 (Figure 2F), probably due to the larger number of lysine residues in LCMV-GP than in RABV-G. Other single-spanning membrane proteins, SARS-CoV-S and SARS-CoV-2-S, which were MARCH8-sensitive in a dose-dependent manner (Figure 1E), harbor four lysine residues in their cytoplasmic tails (at positions 1227, 1237, 1248, and 1251 in SARS-CoV-S and 1245, 1255, 1266, and 1269 in SARS-CoV-2-S) (Figs. 2C and 2D). When these lysines were mutated to arginines, both S mutant proteins displayed relative resistance to MARCH8 (Figs. 2G and 2H). Immunofluorescence confirmed that MARCH8, which induced lysosomal degradation of wild-type envelopes, was unable to downregulate all of these mutant envelopes from the cell surface (Figs. 2I, 2J, 2K and 2L). These results suggest that MARCH8 targets lysine residues located in the cytoplasmic tails of these viral envelopes, leading to their lysosomal degradation that might result from ubiquitination, as previously observed for VSV-G (Zhang et al., 2020).

**Figure 2.**
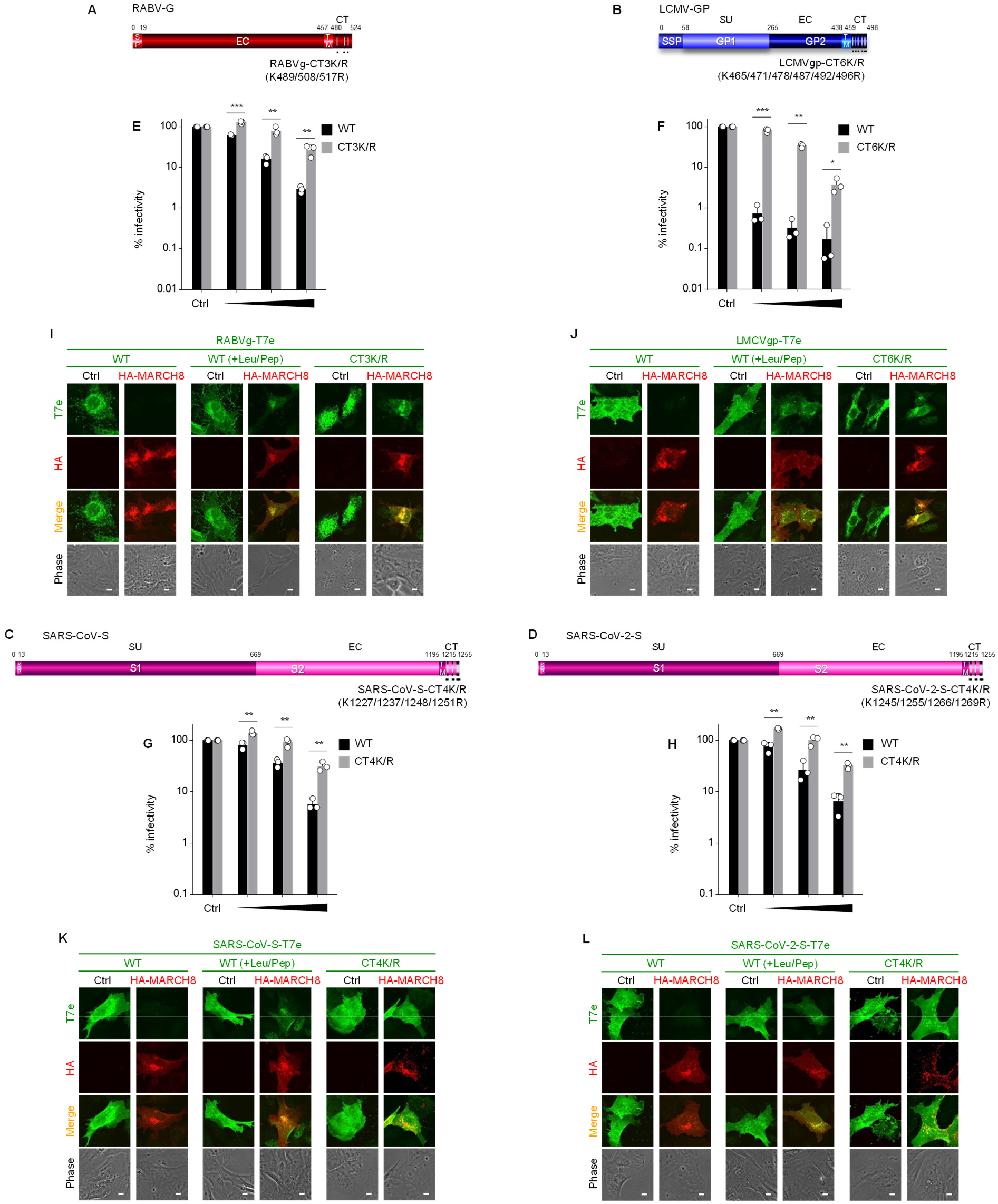
MARCH8 targets lysine residues in the cytoplasmic tail of viral envelopes with single transmembrane domains. **(A-D)** Schematic gene structures of the lysine mutants of **(A)** RABV-G (CT3K/R); **(B)** LCMV-GP (CT-6K/R); **(C)** SARS-CoV-S (CT-4K/R); and **(D)** SARS-CoV-2-S (CT-4K/R). SP, signal peptide; EC, extracellular domain; TM, transmembrane domain; CT, cytoplasmic tail; SU, surface subunit. **(E-H)** MARCH8 resistance is conferred by mutations in cytoplasmic lysine residues of viral envelopes with single transmembrane domains. Infectivity assays were performed as described in Figure 1A except that the maximum amount of MARCH8 plasmid used was 120 ng. Black and gray columns represent the wild-type (WT) and lysine mutants of each viral envelope glycoprotein, respectively. Data from three independent experiments are shown as a percentage of the infectivity of viruses produced in the absence of MARCH8 when the WT protein or its mutant was used (mean ± s.d., *n* = 3 technical replicates). \**p*<0.05, \*\**p*<0.005, \*\*\**p*<0.0005, \*\*\*\**p*<0.0001 compared with the WT using sing two-tailed unpaired *t*-tests. ns, not significant. **(I-L)** Lysine mutants of viral envelopes are resistant to MARCH8-mediated lysosomal degradation. Shown are immunofluorescence-based analyses of the expression of either the T7 epitope (T7e)-tagged WT or K/R mutant of **(I)** RABV-G; **(J)** LCMV-GP; **(K)** SARS-CoV-S; and **(L)** SARS-CoV-2-S with or without MARCH8 in transfected HOS cells. All WT viral envelopes were rescued from MARCH8-induced degradation in the presence of lysosomal protease inhibitors (+Leu/Pep), as shown in each of the middle panels. Scale bars, 10 μm.

### MARCH8 targets a cytoplasmic lysine residue in alphavirus E2 proteins

In the case of CHIKV and RRV polyproteins (E3-E2-6K-E1), cytoplasmic lysines are located in the 6K proteins (Figs. 3A and 3C); therefore, we introduced point mutations into the residues of these proteins and then performed infectivity assays. However, these mutants were found to still be sensitive to MARCH8 at exactly the same levels as the WT envelope (Figs. 3E and 3G), indicating that MARCH8 does not target the lysine residues in the 6K proteins of alphavirus envelope glycoproteins. After being proteolytically cleaved from 6K, the second transmembrane domain of the alphavirus E2 protein is released from the membrane and translocates along with its short ectodomain into the cytoplasm, after which the cleaved E2 protein is associated with E3, 6K, and E1 (Ramsey and Mukhopadhyay, 2017) (Figs. 3B and 3D). Based on this process, we noticed that K422 of CHIKV-E2 could be translocated from the membrane into the cytoplasm after E2 cleavage (Figure 3B), whereas K394 of RRV-E2 was originally exposed in the cytoplasm (Figs. 3C and 3D), both of which were mutated to arginine (K422R and K394R, respectively). The K422R mutation rendered the CHIKV envelope almost completely resistant to MARCH8 (Figure 3F), while the K394 mutation of RRV showed a modest but noticeable effect on MARCH8-mediated inhibition (Figure 3H). The results were consistent with those of immunofluorescence showing that MARCH8-mediated lysosomal degradation observed in the WT envelope (which is C-terminally tagged with the T7 epitope (T7e)) was rescued in the E2-lysine mutants but not in the 6K mutants (Figs. 3I and 3J), suggesting that these two viral envelopes are downregulated by MARCH8 in a manner dependent on E2 domain-specific ubiquitination.

**Figure 3.**
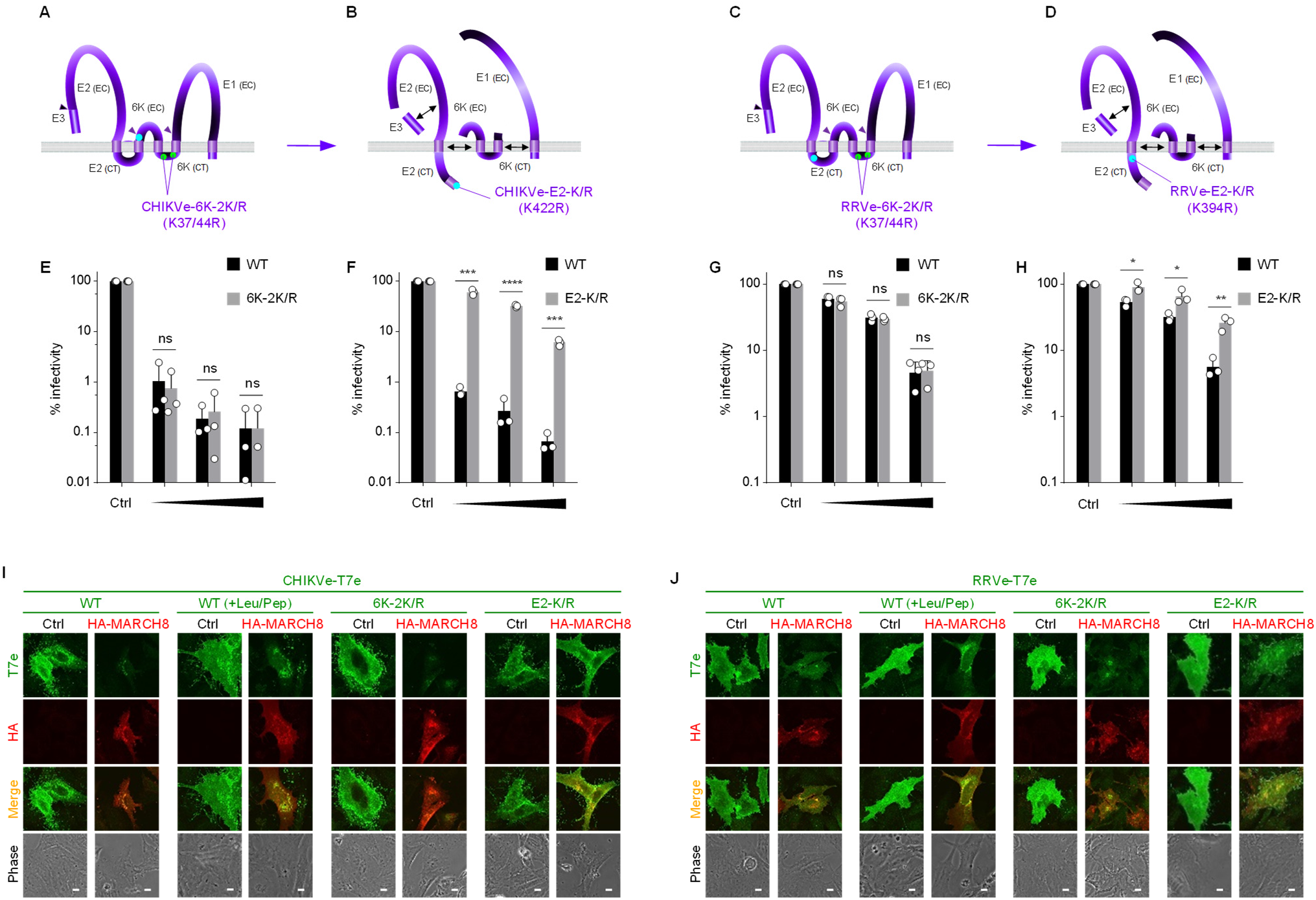
MARCH8 targets a cytoplasmic lysine residue in E2 but not in 6K of alphaviruses. **(A-D)** Schematic representation of topological structures illustrating the multiple-spanning membrane proteins of CHIKV E3-E2-6K-E1 *(A, B)* and RRV E3-E2-6K-E1 *(C, D)*. **(A, C)** In the precleavage state, two lysine residues (37K and 44K, shown in light green) of the 6K protein are exposed to the cytoplasm. These residues were mutated to arginine residues, and the resultant mutants were designated CHIKVe-6K-2K/R *(A)* and RRVe-6K-2K/R *(C)*. Cleavage sites are shown by arrowheads. EC, extracellular domain; CT, cytoplasmic tail. **(B, D)** In the postcleavage state, a lysine residue (K422; light blue) at the C-terminal end of the CHIKV E2 protein, which is present extracellularly in the precleavage state, as shown in A, is relocated and exposed to the cytoplasm *(B)*. In contrast, a lysine residue of RRV E2 (K394; light blue) was originally located at the proximal region of the internal membrane *(D)*. Each lysine residue was replaced by an arginine, and the resultant mutants were designated CHIKVe-E2-2K/R *(B)* and RRVe-E2-2K/R *(D)*. Double-headed arrows represent the putative interaction of cleaved viral proteins. **(E-H)** MARCH8 resistance is conferred by the mutation of a cytoplasmic lysine residue of the alphavirus protein E2 *(F, H)* but not 6K *(E, G)*. Infectivity assays were performed as described in Figure 1A except that the maximum amount of MARCH8 plasmid used was 120 ng. Black and gray columns represent the WT and K/R mutants of alphaviruses, respectively. Data from three independent experiments are shown as a percentage of the infectivity of viruses produced in the absence of MARCH8 when the WT protein or its mutant was used (mean ± s.d., *n* = 3 technical replicates). \**p*<0.05, \*\**p*<0.005, \*\*\**p*<0.0005, \*\*\*\**p*<0.0001 compared with the Ctrl using sing two-tailed unpaired *t*-tests. ns, not significant. **(I, J)** Lysine mutants of alphavirus E2 but not 6K are resistant to MARCH8-mediated lysosomal degradation. Shown are immunofluorescence-based analyses of the expression of either the T7e-tagged WT or K/R mutants of CHIKV-E3-E2-6K-E1 *(I)* and RRV-E3-E2-6K-E1 *(J)* with or without MARCH8 in transfected HOS cells. Both WT viral envelopes were rescued from MARCH8-induced degradation in the presence of lysosomal protease inhibitors (+Leu/Pep). Scale bars, 10 μm.

### MARCH8 ubiquitinates lysine residue(s) at the CT domains of RABV-G and CHIKV-E2

To demonstrate whether the aforementioned cytoplasmic lysine residues of MARCH8-sensitive viral envelope glycoproteins are targets for MARCH8-mediated ubiquitination, we performed immunoprecipitation /Western-based ubiquitination assays using cells coexpressing either control or MARCH8, together with either the RABV-G (which is C-terminally T7e-tagged) or the CHIKV envelope glycoprotein (in which a T7e-tag is inserted immediately downstream of a furin-cleavage site between E3 and E2, resulting in an N-terminally T7e-tagged E2). In cells coexpressing MARCH8, both RABV-G (Figure 4A) and CHIKV-E2 (Figure 4B) were efficiently ubiquitinated, whereas their lysine mutants (CT3K/R and E2-K/R, respectively) did not undergo MARCH8-mediated ubiquitination, as expected, suggesting that lysine residue(s) at either positions 489, 508, and 517 in the CT domain of RABV-G or position 422 in E2 of CHIKV are specifically ubiquitinated by MARCH8. Taken together, these findings indicate that MARCH8 ubiquitinates cytoplasmic lysine residues of a variety of viral envelope glycoproteins and that this ubiquitination leads to lysosomal degradation.

**Figure 4.**
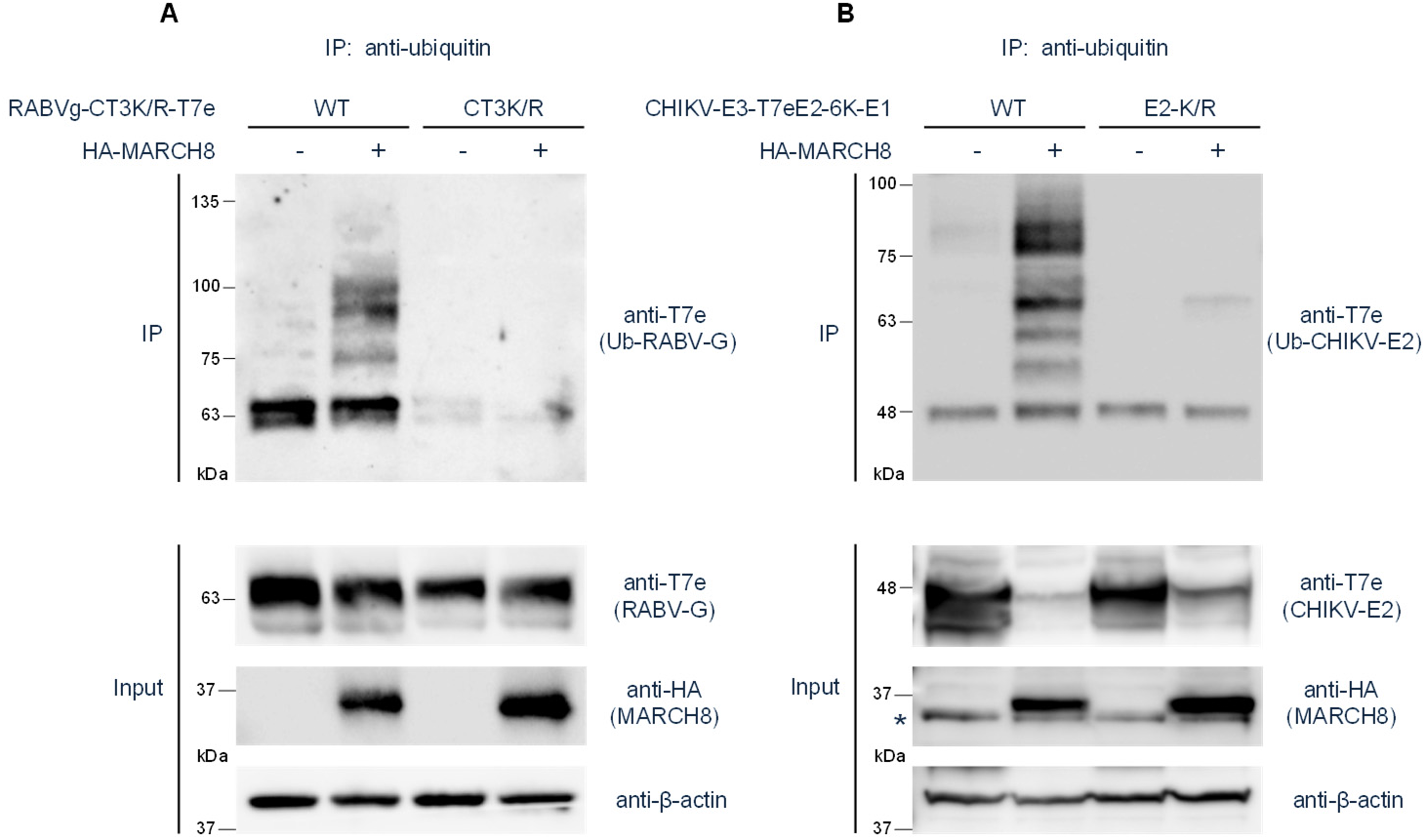
Lysine residue(s) at the CT domains of RABV-G and CHIKV E2 are ubiquitinated by MARCH8. **(A)** The ubiquitination of the WT or CT-lysine mutant of RABV-G tagged with T7e in cells expressing control or HA-tagged MARCH8 was examined by the immunoprecipitation (IP) of ubiquitinated proteins with an anti-ubiquitin antibody, followed by immunoblotting with an anti-T7e antibody. Aliquots of the cell lysates (Input) were also analyzed by immunoblotting for RABV-G (upper), MARCH8 (middle), and β-actin (lower). **(B)** The ubiquitination of the WT or E2-lysine mutant of CHIKV-E3-E2-6K-E1 was similarly investigated (upper). Lysates were also analyzed by immunoblotting for CHIKV-E2 (upper), MARCH8 (middle), and β-actin (lower). Asterisk (middle) indicates non-specific bands.

### The tyrosine motif of MARCH8 is important for its antiviral activity against some viruses

The aforementioned MARCH8-sensitive viral envelope glycoproteins could be considered ubiquitination-sensitive proteins due to the observation of lysine-dependent degradation by MARCH8. Therefore, we examined whether these viral proteins could undergo MARCH8 tyrosine motif-dependent downregulation, as previously observed in HIV-1 Env. As expected, infection by viruses pseudotyped with LCMV-GP or CHIKV-E3-E2-6K-E1, which are highly sensitive to MARCH8 (Figs. 1C and 1E), was still inhibited by the MARCH8 AxxL mutant (Fig. S1), as observed in VSV-G that is extremely MARCH8-sensitive (Tada et al., 2015; Zhang et al., 2020). Conversely, RABV-G, SARS-CoV-S, SARS-CoV-2-S, and RRV-E3-E2-6K-E1, whose functions were dose-dependently abrogated by MARCH8, were almost completely resistant to the AxxL mutant, respectively (Figure S1), suggesting that the inhibition of these viral envelopes is simultaneously regulated by two mechanisms of MARCH8, which are both ubiquitination- and tyrosine motif-dependent downregulations.

### MARCH1 and MARCH2 differentially target viral envelope proteins

We have recently reported that among MARCH family members, MARCH1 and MARCH2 are also antiviral MARCH proteins that inhibit HIV-1 infection (Zhang et al., 2019). Therefore, we examined whether these two MARCH proteins could similarly target various viral envelope glycoproteins, as observed in MARCH8 in this study. Both MARCH1 and MARCH2 were found to be able to differentially inhibit infection by pseudoviruses harboring RABV-G, LCMV-GP, CHIKV and RRV envelope glycoproteins but not the S proteins of SARS-CoV and SARS-CoV-2 although MARCH8 showed the highest inhibitory activity against all viral envelopes tested (Figure S2).

### Antiviral activity of MARCH8 is reproducible in infection with an RABV strain

Finally, because all the aforementioned experiments were performed using pseudotyped viruses, we conducted whole-virus-based infection experiments to verify the inhibitory effects of MARCH8 on not only pseudotyped viruses but also whole viruses. We prepared cells stably expressing MARCH8, used the cells for infection with whole RABV, and then performed immunofluorescence assays using antibodies against two different RABV proteins, phosphoprotein P and envelope glycoprotein G (Figure 5A). As expected, RABV-G protein was completely lost from the surface of MARCH8-expressing cells infected with rabies viruses, whereas the expression of RABV-P was not affected by MARCH8 expression (Figure 5B). Thus, the aforementioned results obtained in pseudotyped viruses were demonstrated to be reproducible in whole-virus infection and were specific for MARCH8-mediated inhibition.

**Figure 5.**
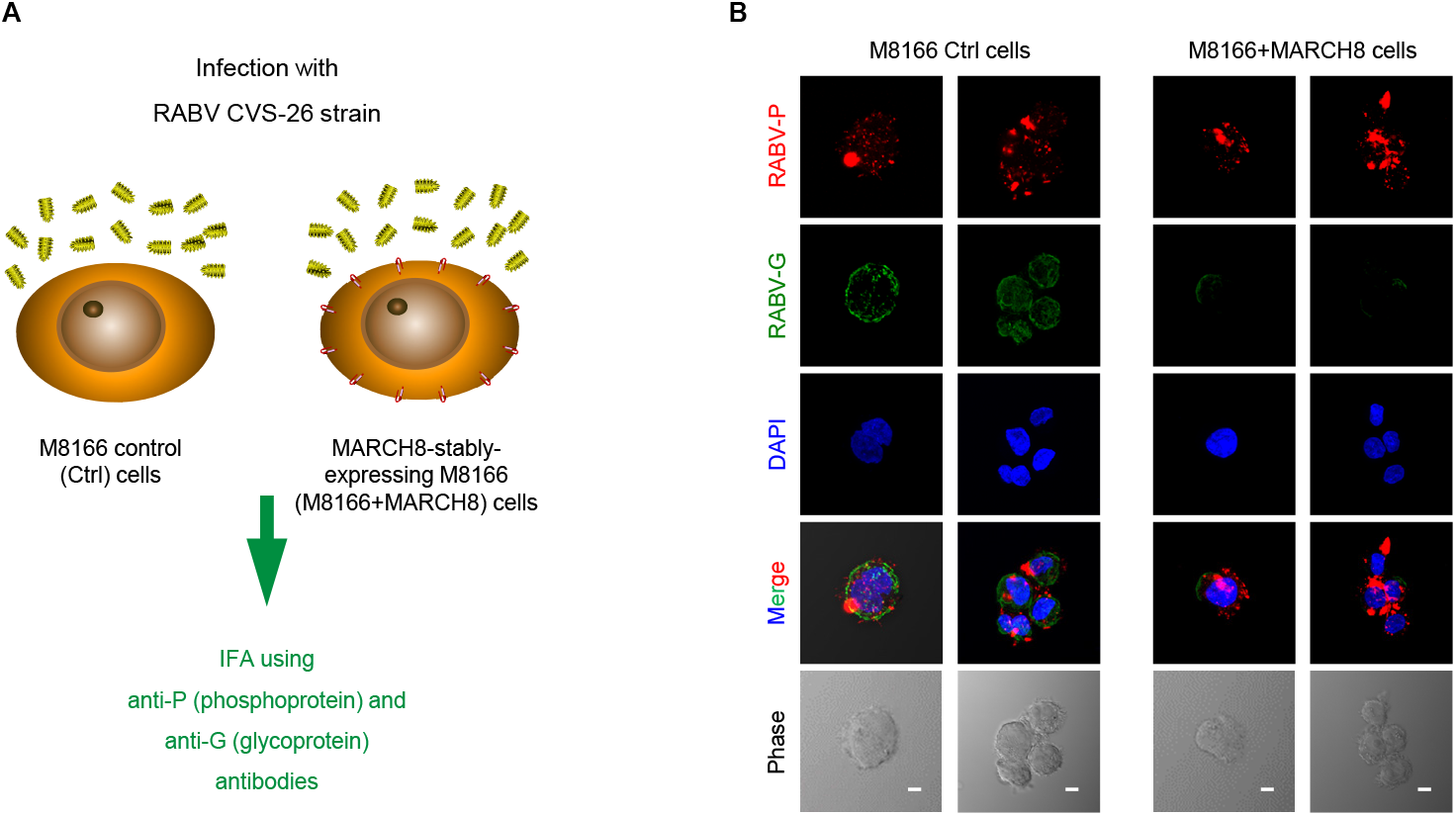
MARCH8-mediated inhibition against pseudoviruses is reproducible in whole-virus infection. **(A)** Schematic flowchart of the infection of MARCH8-stably expressing M8166+MARCH8 cells with the RABV CVS-26 strain. **(B)** MARCH8 downregulates the G protein of the RABV CVS-26 strain. Shown are immunofluorescence images of the cell-surface expression of RABV-G and the intracellular expression of RABV phosphoprotein P (RABV-P) in M8166 control (Ctrl) cells (left panels) and M8166+MARCH8 cells (right panels). Scale bars, 10 μm.

## Discussion

In this study, we show that MARCH8 can target a broad spectrum of viral envelope glycoproteins. Among viral envelopes that have single-spanning membrane regions, MARCH8-sensitive proteins include those derived from not only retroviruses and a rhabdovirus VSV, as we previously reported (Tada et al., 2015), but also another rhabdovirus RABV, an arenavirus LCMV, and the coronaviruses SARS-CoV/SARS-CoV-2. Independent of virus family, the levels of MARCH8 sensitivity might be determined, to some extent, by the number of lysine residues in their cytoplasmic tails (Figure S3) that could be ubiquitinated by MARCH8 under the condition that lysines are optimally exposed at which the ubiquitin ligase is readily accessible. Indeed, among single-spanning viral envelopes, RABV-G and pathogenic coronavirus S proteins (from SARS-CoV and SARS-CoV-2), whose cytoplasmic tails harbors three and four lysines out of 44 and 39 amino acids, respectively (Figure S3), are moderately MARCH8-sensitive (Figure 1B). In contrast, VSV-G and LCMV-GP, whose cytoplasmic tails are lysine-rich (5 out of 29 and 6 out of 39 amino acids, respectively; Figure S3), are highly susceptible to MARCH8 (Tada et al., 2015) and Figure 1C) but are, in turn, resistant to this host antiviral protein when their lysine residues are mutated to arginines. It should be noted that HIV-1 Env carries only two lysines out of 151 amino acids in the cytoplasmic tail but is still MARCH8-sensitive when these lysines are mutated to arginines (Zhang et al., 2020).

Other viral envelopes sensitive to MARCH8 are those derived from alphaviruses (CHIKV and RRV), whose envelope glycoproteins are initially expressed as polyproteins with multiple transmembrane domains. These polyproteins are proteolytically cleaved, resulting in single-spanning membrane proteins formed with E2-E1 heterodimers (Ramsey and Mukhopadhyay, 2017). In the case of CHIKV-E3-E2-6K-E1, which is extremely sensitive to MARCH8, a lysine mutation in E2 conferred strong resistance to MARCH8, suggesting that the lysine residue at the C-terminal edge of cleaved E2 (Figure 3B) can easily access MARCH8 in the cytoplasm and is therefore critical for MARCH8 sensitivity. Based on the fact that only a moderate effect was observed in a mutation of the lysine located in the membrane-proximal position of the RRV-E2 cytoplasmic tail (Figure 3D), it is likely that not only the number but also the location of the lysine residues, which possibly affects the accessibility of MARCH8, may also determine the sensitivity to MARCH8.

Unexpectedly, in the absence of MARCH8, the luciferase values before individual normalization showed that the infectivity of the aforementioned cytoplasmic lysine mutants was higher than that of their wild-type envelopes to varying degrees (1.5-3-fold higher in those of RABV, LCMV, SARS-CoV, and CHIKV; 5-fold higher in that of SARS-CoV-2), except for that of the RRV-6K mutant (Figure S4). Because our previous findings demonstrated that the endogenous expression of MARCH8 in 293T cells (also used for virus production in this study) is extremely low (Tada et al., 2015), it is unlikely that these viral envelopes harboring lysine mutations successfully evaded inhibition by the low levels of MARCH8 in this cell line. Therefore, we postulate that unknown E3 ubiquitin ligase(s) endogenously expressed in 293T cells might be partially involved in targeting the lysine residues in the cytoplasmic tails of viral envelopes, although this hypothesis needs to be verified by further experiments.

Among the MARCH8-sensitive viral envelopes tested in this study, phenotypes of viral envelopes are divided into two groups: (i) highly MARCH8-sensitive in the WT envelope and resistant in its lysine mutant (VSV-G, LCMV-GP, and CHIKV-E3-E2-6K-E1); and (ii) dose-dependently MARCH8-sensitive in the WT envelope and partially resistant in its lysine mutant (RABV-G, SARS-CoV-S, SARS-CoV-2-S, and RRV-E3-E2-6K-E1). In both types, MARCH8 clearly targets the cytoplasmic lysine residues of the viral envelopes. Interestingly, the difference in WT MARCH8 sensitivity paralleled that in the resistance to the MARCH8 Yxxϕ mutant; i.e., whereas viral envelopes in the case of (i) were still sensitive to the Yxxϕ mutant, those in the cases of (ii) were resistant to this mutant. These results suggest that MARCH8 downregulates (a) VSV-G, LCMV-GP, and CHIKV-E3-E2-6K-E1 in a ubiquitination-dependent pathway (Figure S5, left) and (b) RABV-G, SARS-CoV-S, SARS-CoV-2-S, and RRV-E3-E2-6K-E1 in both ubiquitination- and Yxxϕ motif-dependent pathways (Figure S5, middle). As an exceptional case compared with all other viral envelopes tested here, it is intriguing that MARCH8-induced downregulation of HIV-1 Env is ubiquitination-independent and Yxxϕ motif-dependent, as we recently reported (Zhang et al., 2020) (Figure S5, right), and the ubiquitination independency is indeed consistent with a very recent paper (Lun et al., 2021) showing that MARCH8-mediated restriction of HIV-1 Env was independent of the presence of its cytoplasmic tail, which carries two lysine residues.

During the preparation of this manuscript, three reports, including the above (Lun et al., 2021; Umthong et al., 2021; Yu et al., 2020), showed that in addition to HIV-1 Env and VSV-G, MARCH8 has broad antiviral functions targeting viral glycoproteins from influenza virus, Ebola virus, murine leukemia virus, and Nipah virus, along with those from SARS-CoV-2, LCMV, and CHIKV, as observed in this study. In the case of SARS-CoV-2, Lun *et a*l. showed that its lysine residues do not affect sensitivity to MARCH8, probably due to the results obtained from single-dose experiments (Lun et al., 2021). Nevertheless, the current consensus on the function of MARCH8 from these multiple studies, including ours, is that MARCH8 has a broad antiviral spectrum against various viral envelope glycoproteins, as we originally expected, and here, we conclude that MARCH8 targets the cytoplasmic lysine residues of these viral envelopes. Further investigations will clarify the host defense mechanisms of this protein in more detail.

## Materials and methods

### DNA constructs

The envelope glycoprotein (Env)-deficient HiBiT-tagged HIV-1 proviral indicator construct pNL-Luc2-IN/HiBiT-E(-)Fin, the HIV-1 Gag-Pol expression plasmid pC-GagPol-RRE, the VSV-G expression plasmid pC-VSVg, the severe acute respiratory syndrome coronavirus (SARS-CoV) spike (S) protein expression plasmid pC-SARS-S, the SARS-CoV-2-S protein expression plasmid pC-SARS2-S, the HIV-1 Rev expression plasmid pCa-Rev, the MARCH8 expression plasmid pC-MARCH8, pC-HA-MARCH8, and the tyrosine motif-mutant pC-MARCH8-AxxL, the ACE2 expression plasmid pC-ACE2 and the TMPRSS2 expression plasmid pC-TMPRSS2, have previously been described elsewhere (Iwabu et al., 2009; Ozono et al., 2021; Ozono et al., 2020; Tada et al., 2015). The plasmid pC-LCMVgp expressing the viral envelope derived from lymphocytic choriomeningitis virus (LCMV) glycoprotein (gp) was created by inserting PCR-amplified and BsiWI/XhoI-digested LCMV-GP fragments (PCR-amplified from pCI-LCMV-GP (Reignier et al., 2006)) into the Acc65I/XhoI-digested pCAGGS mammalian expression plasmid. The plasmid pC-CHIKVe expressing alphavirus E3-E2-6K-E1 polyprotein derived from Chikungunya virus (CHIKV) was created by inserting the PCR-amplified and Acc65I/XhoI-digested E3-E2-6K-E1 fragments (PCR-amplified from codon-optimized CHIKV’s C-E3-E2-6K-E1 plasmid (Kishishita et al., 2013)) into the corresponding site of pCAGGS. Another E3-E2-6K-E1 expression plasmid, pC-RRVe, derived from the Ross River virus (RRV) T48 strain, was created by inserting the PCR-amplified and EcoRV/NotI-digested E3-E2-6K-E1 fragments (PCR-amplified from pRRV-E2E1 (Sharkey et al., 2001)) into the corresponding site of pCAGGS. The plasmid pC-RABVg expressing the glycoprotein (G) of rabies virus (RABV) was created by inserting the RT-PCR-amplified and Acc65I/XmaI-digested G fragments (RT-PCR-amplified from total RNA from CVS rabies strain-infected cells (Fernandes et al., 1964) into the corresponding site of pCAGGS. The RABV-G or LCMV-GP mutant (pC-RABVg-CT3K/R, or pC-LCMVgp-CT6K/R), in which cytoplasmic lysine residues at positions 489/508/517 or 465/471/478/487/492/496 were mutated to arginine residues, was created by inserting overlapping PCR fragments into correspondingly digested pC-RABVg or pC-LCMVgp, respectively. Mutants of SARS-CoV-S and SARS-CoV-2-S (pC-SARS-S-CT4K/R, or pC-SARS2-S-CT4K/R), in which cytoplasmic lysine residues at positions 1227/1237/1248/1251 and 1245/1255/1266/1269 were mutated to arginine residues, were created by inserting overlapping PCR fragments into correspondingly digested pC-SARS-S and pC-SARS2-S, respectively. The CHIKV and RRV 6K mutants (pC-CHIKV-6K-2K/R and pC-RRV-6K-2K/R), in which the 6K region’s cytoplasmic lysine residues at positions 37/44 were mutated to arginine residues, were created by inserting overlapping PCR fragments into correspondingly digested pC-CHIKVe and pC-RRVe, respectively. Similarly, the E2 mutants of CHIKV and RRV (pC-CHIKVe-E2-K/R or pC-RRVe-E2-K/R), in which E2’s C-terminal lysine residues at positions 422 and 394 were mutated to arginine residues, were created by inserting overlapping PCR fragments into correspondingly digested pCAGGS. The C-terminally T7-epitope–tagged wild-type and its mutant expression plasmids of RABV-G (pC-RABVg-T7e and pC-RABVg-CT3K/R-T7e), LCMV-GP (pC-LCMVgp-T7e and pC-LCMVgp-CT6K/R-T7e), SARS-CoV-S (pC-SARS-S-T7e and pC-SARS-S-CT4K/R-T7e), SARS-CoV-2-S (pC-SARS2-S-T7e and pC-SARS2-S-CT4K/R-T7e), CHIKVe3-E2-6K-E1 (pC-CHIKVe-T7e, pC-CHIKVe-6K-2K/R-T7e, and pC-CHIKVe-E2-K/R-T7e), and RRVe3-E2-6K-E1 (pC-RRVe-T7e, pC-RRVe-6K-2K/R-T7e, and pC-RRVe-E2-K/R-T7e) were created by replacing the VSV-G gene of pC-VSVg-T7e (Zhang et al., 2020) with the corresponding PCR-amplified fragments. To detect ubiquitinated CHIKV-E2 proteins by immunoprecipitation assays, WT or E2-K/R mutant plasmids expressing CHIKV E3-E2-6K-E1, in which the T7 epitope-tag is inserted immediately downstream of a furin-cleavage site between E3 and E2, were created by inserting overlapping PCR fragments that harbor a T7e sequence into correspondingly digested pCAGGS and were designated pC-CHIKVe-T7eE2, pC-CHIKVe-T7eE2-K/R, pC-RRVe-T7eE2, and pC-RRVe-T7eE2-K/R, respectively. All constructs were verified by a DNA sequencing service (FASMAC).

### Cell maintenance

293T, HeLa, NIH3T3, HOS, and M8166+MARCH8 (Tada et al., 2015) cells were maintained under standard conditions. Cells were originally obtained from ATCC and routinely tested negative for mycoplasma contamination (PCR Mycoplasma Detection Set, Takara).

### Virion infectivity assays

To prepare various viral envelope-pseudotyped HIV-1 luciferase reporter viruses, 1.1 × 10^5^ 293T cells were cotransfected with increasing amounts of the MARCH8 expression plasmid, 20 ng of viral envelope expression plasmid (pC-VSVg, pC-RABVg, pC-LCMVgp, pC-SARS-S, pC-SARS2-S, pC-CHIKVe, pC-RRVe, and their mutants), 500 ng of pNL-Luc2-IN/HiBiT-E(-)Fin, and an empty vector up to 1 μg of total DNA using FuGENE 6 (Promega). Sixteen hours later, transfected cells were washed with phosphate-buffered saline, and 1 ml of fresh complete medium was added. After 24 h, the supernatants were harvested and treated with 37.5 U/ml DNase I (Roche) at 37 °C for 30 min. Viral supernatants were measured by the HiBiT assay, as previously described (Ozono et al., 2020). Briefly, a standard virus stock with known levels of p24 antigen was serially diluted. Either the standards or viral supernatants containing pseudotyped viruses (25 μl) and LgBiT Protein (1:100)/HiBiT Lytic Substrate (1:50) in Nano-Glo HiBiT Lytic Buffer (25 μl) (Nano-Glo HiBiT Lytic Detection System; Promega) were mixed and incubated for ten minutes at room temperature according to a modified version of the manufacturer’s instructions. HiBiT-based luciferase activity in viral supernatants was determined with a Centro LB960 luminometer (Berthold) and was translated into p24 antigen levels. To determine viral infectivity, 1 × 10^4^ NIH3T3 cells or HeLa cells were incubated with 1 ng of p24 antigen of either arenavirus virus envelope (LCMV-GP)- or alphavirus E3-E2-6K-E1 envelope (derived from CHIKV or RRV)-pseudotyped HIV-1 luciferase reporter viruses, respectively.

Additionally, 2.2 × 10^4^ 293T cells were incubated with 1 ng of p24 antigen of HIV-1 luciferase reporter viruses pseudotyped with VSV-G or RABV-G. Alternatively, 2.2 × 10^4^ 293T cells transiently coexpressing ACE2 and TMPRSS2 (using pC-ACE2 and pC-TMPRSS2) were incubated with 1 ng of p24 antigen of either SARS-CoV-S- or SARS-CoV-2-S-pseudotyped luciferase reporter lentiviruses. After 48 h, cells were lysed in 100 μl of One-Glo Luciferase Assay Reagent (Promega). The firefly luciferase activity was determined with a Centro LB960 (Berthold) luminometer.

### Immunofluorescence microscopy

HOS cells were plated on 13-mm coverslips, cotransfected with 0.5 μg of pC-xx-T7e (xx: RABV-G, RABV-G-CT3K/R, LCMVgp, LCMVgp-CT6K/R, SARS-S, SARS-S-CT4K/R, SARS2-S, SARS2-S-CT4K/R, CHIKVe, CHIKVe-6K-2K/R, CHIKVe-E2-K/R, RRVe, RRVe-6K-2K/R, or RRVe-E2-K/R), 0.1 μg of pC-GagPol-RRE, 0.05 μg of pCa-Rev, and 0.3 μg of either the HA-MARCH8 expression plasmid or an empty control using FuGENE6, and cultured for 24 h. To examine whether WT envelope glycoproteins are lysosomally degraded by MARCH8, the transfected cells were cultured in the presence of lysosome protease inhibitors (40 μM of leupeptin and pepstatin A) for 14 h. For the total staining of both Env-T7es and HA-MARCH8, cells were fixed with 4% paraformaldehyde for 30 min on ice and permeabilized with 0.05% saponin. Fixed cells were incubated with anti-T7e epitope mouse monoclonal antibodies (Novagen, 69522-4) and anti-HA rabbit polyclonal antibodies (Sigma Aldrich, H6908). The secondary antibodies Alexa 488 donkey anti-mouse IgG (Molecular Probes, A-21202) and Alexa 568 donkey anti-rabbit IgG (Molecular Probes, A-10042) were used for double staining. Confocal images were obtained with a FluoView FV10i automated confocal laser-scanning microscope (Olympus). For whole-virus infection of cells stably expressing MARCH8, M8166+MARCH8 cells were plated on CELLview dishes (35-mm, 4-compartment cell culture dishes with a glass bottom; Greiner Bio-One), infected with the RABV CVS-26 strain (Hamamoto et al., 2015) at a multiplicity of infection of 0.5, and cultured for 24 h. To detect cell-surface RABV-G protein, cells were fixed with 10% formalin neutral buffer solution, pH 7.4 (Wako Pure Chemical, Osaka, Japan), at room temperature for 30 min. Fixed cells were incubated with an anti-RABV-G mouse monoclonal #7-1-9 (0.4 mg/ml), kindly provided by Akihiko Kawai (Research Institute for Production Development, Kyoto, Japan) (Irie and Kawai, 2002). To detect intracellular RABV-P protein, cells were permeabilized with 0.2% Triton X-100 in PBS for 5 min at room temperature and incubated with anti-RABV-P rabbit polyclonal antibody (Tobiume et al., 2009). The secondary antibodies Alexa 488 donkey anti-mouse IgG and Alexa 568 donkey anti-rabbit IgG were used for staining. Nuclei were visualized by DAPI staining. Confocal images were obtained with an FV1000 confocal laser scanning microscope (Olympus).

### Ubiquitination assays

293T cells (5 × 10^5^) were cotransfected with 0.8 μg of pC-RABVg-T7e, pC-RABVg-CT3K/R-T7e, pC-CHIKVe-T7eE2, or pC-CHIKVe-T7eE2-K/R; and 0.2 μg of pC-HA-MARCH8 or an empty control. After 48 h, cells were lysed in TBS-T buffer (50 mM Tris-HCl buffer (pH 7.5), 0.15 M NaCl, 1% Triton X-100, and 0.5% deoxycholic acid) containing protease inhibitor cocktail and 10 mM N-ethylmaleimide as an inhibitor of deubiquitination enzymes. The mixture was centrifuged at 21,500 × g for 15 min, and the supernatant was used as total cell lysate for immunoblotting or immunoprecipitation. Fifty microliters of Protein A-coupled Sepharose 4B (GE Healthcare, 17-0780-01) was preincubated for 2 h at 4 °C with 4 μg of anti-ubiquitin mouse monoclonal antibodies (Clone FK2, Cayman, 14220). Total cell lysate was incubated with antibody-coupled Sepharose for 20 h at 4 °C. The Sepharose was washed three times with TBS-T buffer and one time with PBS before the immunoprecipitated proteins were eluted with SDS sample buffer. To evaluate ubiquitination states, proteins immunoprecipitated with anti-ubiquitin mouse antibodies were subjected to Western blotting with anti-T7e rabbit antibodies. Total cell lysate was also subjected to immunoblotting with anti-T7e rabbit antibodies and anti-HA rabbit polyclonal antibodies (Sigma Aldrich, H6908) to evaluate the expression levels of Env-T7es and HA-MARCH8. Immunoreactive bands were detected using an ECL detection kit (ATTO, EzWestLumi plus, WSE-7120) with a ChemiDoc imaging system (BIO-RAD).

### Statistical analyses

Column graphs that combine bars and individual data points were created with GraphPad Prism version 9.10. *P*-values generated from one-way analysis of variance and Dunnett’s multiple comparison tests for data represented in Figures 1 (B, C, D, and E), S1, and S2, and from two-tailed unpaired t-tests for data represented in Figures 2 (I, J, K, and L), and 3 (E, F, G, and, H).

## Acknowledgments

This work was supported by grants from the Japan Society for the Promotion of Science (KAKENHI, 18K07156 and 21K07060) to K. T.

## Author contributions

Y.Z., S.O., T.T., M.T., M.K., H.F. and K.T. performed experiments and analyzed the data. Y.Z., T.T., H.F. and K.T. discussed the data. M.T., M.K. and S.K. provided reagents. K.T. conceived the study, supervised the work and wrote the paper. All authors read and approved the final manuscript.

## Supplementary information

**Figure S1.**
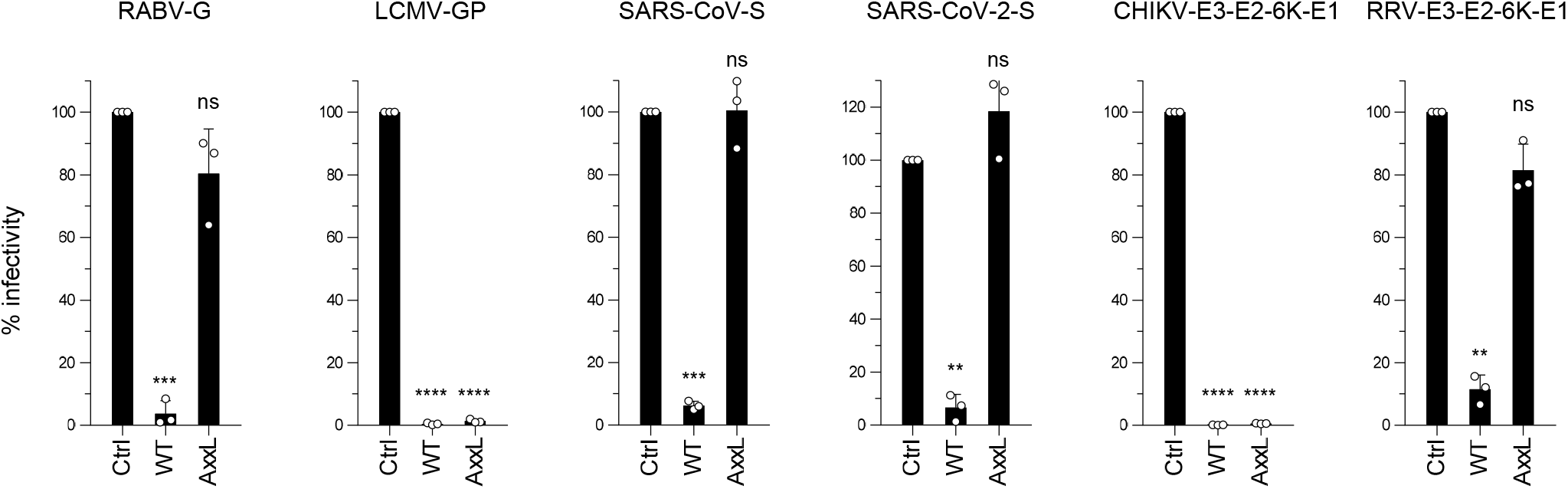
The tyrosine motif of MARCH8 partially mediates the downregulation of RABV-G, SARS-CoV-S, SARS-CoV-2-S, and RRV-E3-E2-6K-E1 but not LCMV-GP or CHIKV-E3-E2-6K-E1. Infectivity of viruses prepared from 293T cells cotransfected with Env-defective HiBiT-tagged HIV-1 luciferase (luc) reporter proviral DNA clone pNL-Luc2-IN/HiBiT-E(-) and a control (Ctrl), WT, or ^222^AxxL^225^ MARCH8 plasmid, together with an RABV-G, LCMV-GP, SARS-CoV-S, SARS-CoV-2-S, CHIKV-E3-E2-6K-E1, or RABV-E3-E2-6K-E1 expression plasmid. Data are shown as a percentage of the viral infectivity in the absence of MARCH8 (mean + s.d. from three independent experiments). \**p*<0.05, \*\**p*<0.005, \*\*\**p*<0.0005, \*\*\*\**p*<0.0001 compared with the Ctrl using one-way analysis of variance and Dunnett’s multiple comparison tests. ns, not significant.

**Figure S2.**
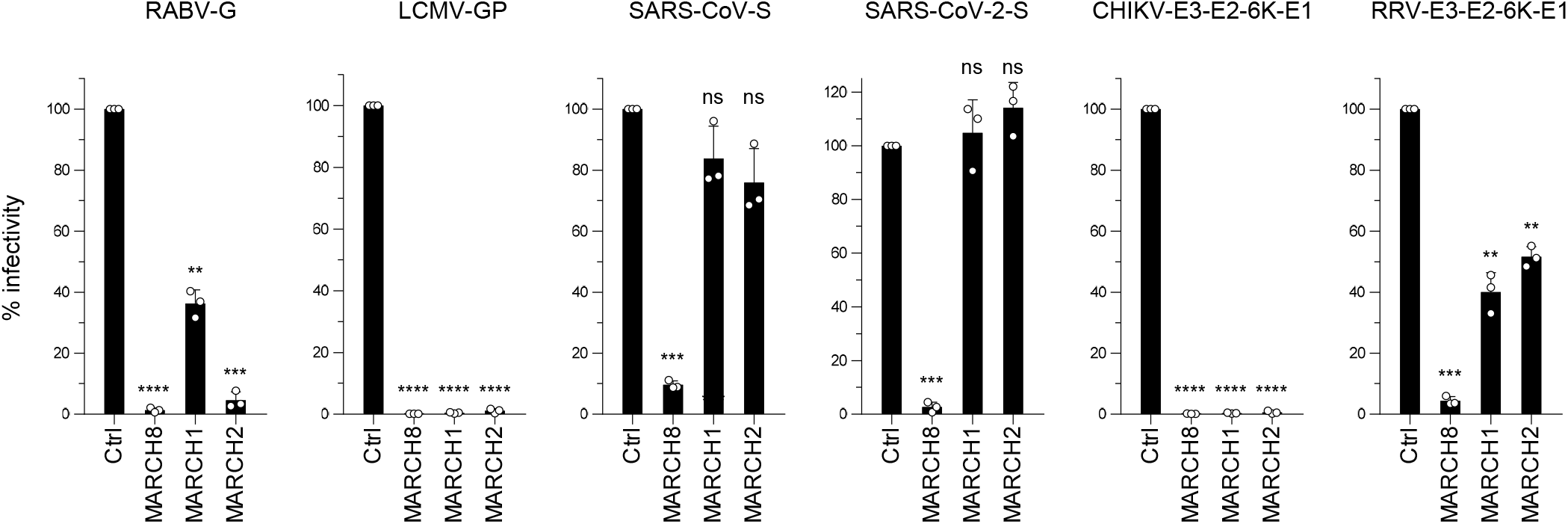
MARCH1 and MARCH2 differentially target a variety of viral envelope proteins. Infectivity of viruses prepared from 293T cells cotransfected with Env-defective HiBiT-tagged HIV-1 luciferase (luc) reporter proviral DNA clone pNL-Luc2-IN/HiBiT-E(-) and a control (Ctrl), MARCH1, MARCH2, or MARCH8 plasmid, together with an RABV-G, LCMV-GP, SARS-CoV-S, SARS-CoV-2-S, CHIKV-E3-E2-6K-E1, or RABV-E3-E2-6K-E1 expression plasmid. Data are shown as a percentage of the viral infectivity in the absence of MARCH8 (mean + s.d. from three independent experiments). \**p*<0.05, \*\**p*<0.005, \*\*\**p*<0.0005, \*\*\*\**p*<0.0001 compared with the Ctrl using one-way analysis of variance and Dunnett’s multiple comparison tests. ns, not significant.

**Figure S3.**
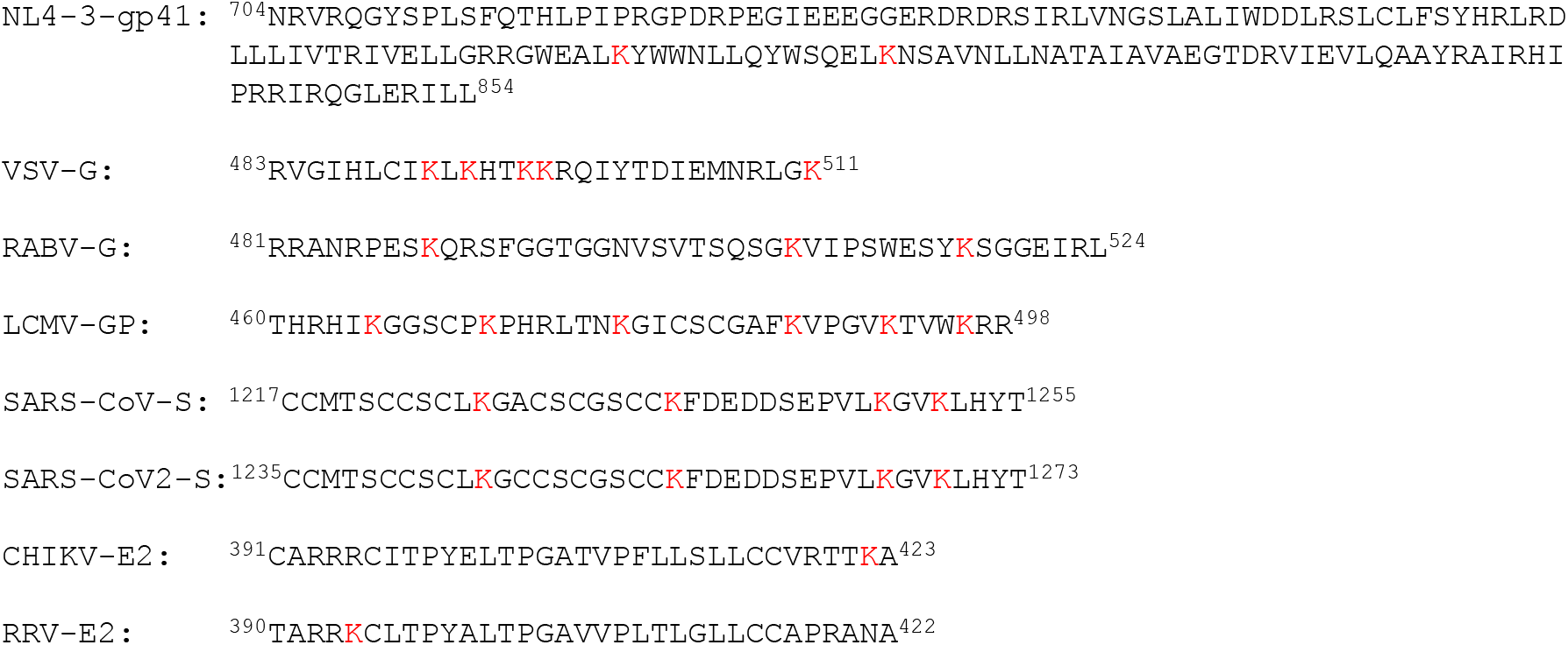
Amino acid sequences of the cytoplasmic tails of the viral envelope glycoproteins tested in our previous and current studies. Among cytoplasmic tail sequences shown above, the alphavirus E2 proteins (CHIKV-E2 and RRV-E2) contain the second transmembrane domains, which are released from the membrane after its cleavage and is then translocated (along with its short ectodomain) into the cytoplasm.

**Figure S4.**
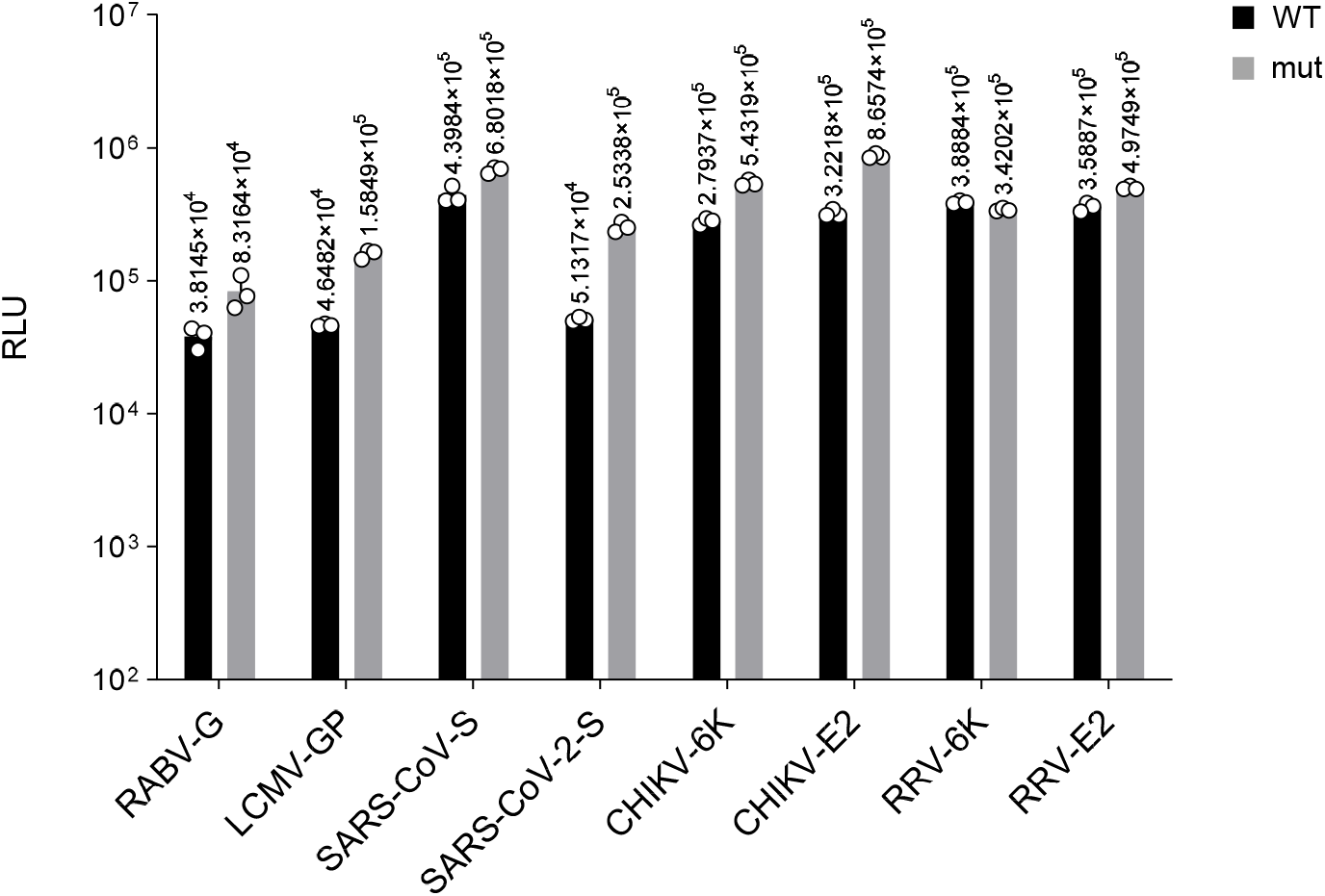
Firefly luciferase values before individual normalization. Shown are the firefly luciferase values obtained in infection of different target cells with the indicated wild-type (WT; black) or mutant (mut; gray) pseudoviruses produced from transfected 293T cells in the absence of MARCH8. Representative data from three independent experiments are shown.

**Figure S5.**
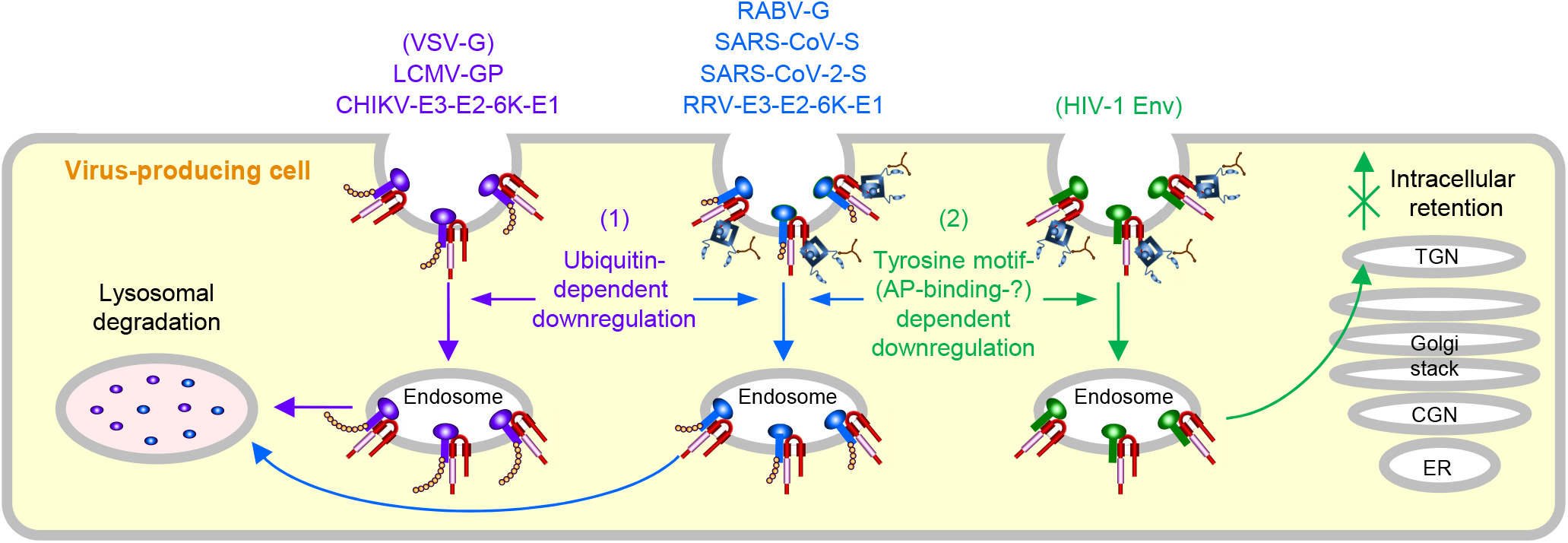
Schematic diagram of different mechanisms by which MARCH8 downregulates viral envelopes. *Left*, MARCH8 (red) downregulates VSV-G, LCMV-GP, and CHIKV-E3-E2-6K-E1 (violet) in a ubiquitin-dependent manner. The RING-CH domain (pink) of MARCH8 mediates ubiquitin conjugation (shown as orange beads) at cytoplasmic lysine residue(s) of VSV-G, LCMV-GP and CHIKV-E2, leading to lysosomal degradation. *Right*, MARCH8 downregulates HIV-1 Env (green) in a tyrosine motif-dependent manner, as previously described (Zhang *et al*. 2020). The tyrosine motif in the C-terminal CT of MARCH8 likely interacts with the adaptor protein μ-subunit (navy) (involving clathrin (brown) if this is the case with μ2 or μ1), resulting in the TGN retention of HIV-1 Env. *Middle*. MARCH8 downregulates RABV-G, SARS-CoV-S, SARS-CoV-2-S, and RRV-E3-E2-6K-E1 (blue) in both ubiquitin- and tyrosine motif-dependent manners, leading to lysosomal degradation. Note that MARCH8-induced downregulation of these viral glycoproteins might not necessarily occur at the plasma membrane. The nucleus and other organelles are not shown. The image was adapted from that of Zhang *et al*. (*eLife*, 9:e57763, 2020).

## References

Baravalle, G., Park, H., McSweeney, M., Ohmura-Hoshino, M., Matsuki, Y., Ishido, S. and Shin, J. S. (2011). Ubiquitination of CD86 is a key mechanism in regulating antigen presentation by dendritic cells. J Immunol 187, 2966–73.

Bartee, E., Eyster, C. A., Viswanathan, K., Mansouri, M., Donaldson, J. G. and Fruh, K. (2010). Membrane-Associated RING-CH proteins associate with Bap31 and target CD81 and CD44 to lysosomes. PLoS One 5, e15132.

Bartee, E., Mansouri, M., Hovey Nerenberg, B. T., Gouveia, K. and Fruh, K. (2004). Downregulation of major histocompatibility complex class I by human ubiquitin ligases related to viral immune evasion proteins. J Virol 78, 1109–20.

Chen, R., Li, M., Zhang, Y., Zhou, Q. and Shu, H. B. (2012). The E3 ubiquitin ligase MARCH8 negatively regulates IL-1beta-induced NF-kappaB activation by targeting the IL1RAP coreceptor for ubiquitination and degradation. Proc Natl Acad Sci U S A 109, 14128–33.

Eyster, C. A., Cole, N. B., Petersen, S., Viswanathan, K., Fruh, K. and Donaldson, J. G. (2011). MARCH ubiquitin ligases alter the itinerary of clathrin-independent cargo from recycling to degradation. Mol Biol Cell 22, 3218–30.

Fernandes, M.V., Wiktor, T.J. and Koprowski, H. (1964). ENDOSYMBIOTIC RELATIONSHIP BETWEEN ANIMAL VIRUSES AND HOST CELLS : A STUDY OF RABIES VIRUS IN TISSUE CULTURE. J Exp Med 120, 1099–1115.

Fujita, H., Iwabu, Y., Tokunaga, K. and Tanaka, Y. (2013). Membrane-associated RING-CH (MARCH) 8 mediates the ubiquitination and lysosomal degradation of the transferrin receptor. J Cell Sci 126, 2798–809.

Funakoshi, Y., Chou, M. M., Kanaho, Y. and Donaldson, J. G. (2014). TRE17/USP6 regulates ubiquitylation and trafficking of cargo proteins that enter cells by clathrin-independent endocytosis. J Cell Sci 127, 4750–61.

Goto, E., Ishido, S., Sato, Y., Ohgimoto, S., Ohgimoto, K., Nagano-Fujii, M. and Hotta, H. (2003). c-MIR, a human E3 ubiquitin ligase, is a functional homolog of herpesvirus proteins MIR1 and MIR2 and has similar activity. J Biol Chem 278, 14657–68.

Hamamoto, N., Uda, A., Tobiume, M., Park, C. H., Noguchi, A., Kaku, Y., Okutani, A., Morikawa, S. and Inoue, S. (2015). Association between RABV G Proteins Transported from the Perinuclear Space to the Cell Surface Membrane and N-Glycosylation of the Sequon Asn(204). Jpn J Infect Dis 68, 387–93.

Irie, T. and Kawai, A. (2002). Studies on the different conditions for rabies virus neutralization by monoclonal antibodies #1-46-12 and #7-1-9. J Gen Virol 83, 3045–3053.

Iwabu, Y., Fujita, H., Kinomoto, M., Kaneko, K., Ishizaka, Y., Tanaka, Y., Sata, T. and Tokunaga, K. (2009). HIV-1 accessory protein Vpu internalizes cell-surface BST-2/tetherin through transmembrane interactions leading to lysosomes. J Biol Chem 284, 35060–72.

Jahnke, M., Trowsdale, J. and Kelly, A. P. (2013). Ubiquitination of HLA-DO by MARCH family E3 ligases. Eur J Immunol 43, 1153–61.

Kishishita, N., Takeda, N., Anuegoonpipat, A. and Anantapreecha, S. (2013). Development of a pseudotyped-lentiviral-vector-based neutralization assay for chikungunya virus infection. J Clin Microbiol 51, 1389–95.

Lun, C. M., Waheed, A. A., Majadly, A., Powell, N. and Freed, E. O. (2021). Mechanism of Viral Glycoprotein Targeting by Membrane-Associated RING-CH Proteins. mBio 12.

Ohmura-Hoshino, M., Matsuki, Y., Aoki, M., Goto, E., Mito, M., Uematsu, M., Kakiuchi, T., Hotta, H. and Ishido, S. (2006). Inhibition of MHC class II expression and immune responses by c-MIR. J Immunol 177, 341–54.

Ozono, S., Zhang, Y., Ode, H., Sano, K., Tan, T. S., Imai, K., Miyoshi, K., Kishigami, S., Ueno, T., Iwatani, Y. et al. (2021). SARS-CoV-2 D614G spike mutation increases entry efficiency with enhanced ACE2-binding affinity. Nat Commun 12, 848.

Ozono, S., Zhang, Y., Tobiume, M., Kishigami, S. and Tokunaga, K. (2020). Super-rapid quantitation of the production of HIV-1 harboring a luminescent peptide tag. J Biol Chem 295, 13023–13030.

Ramsey, J. and Mukhopadhyay, S. (2017). Disentangling the Frames, the State of Research on the Alphavirus 6K and TF Proteins. Viruses 9, 228.

Reignier, T., Oldenburg, J., Noble, B., Lamb, E., Romanowski, V., Buchmeier, M. J. and Cannon, P. M. (2006). Receptor use by pathogenic arenaviruses. Virology 353, 111–120.

Sharkey, C. M., North, C. L., Kuhn, R. J. and Sanders, D. A. (2001). Ross River virus glycoprotein-pseudotyped retroviruses and stable cell lines for their production. J Virol 75, 2653–2659.

Singh, S., Bano, A., Saraya, A., Das, P. and Sharma, R. (2021). iTRAQ-based analysis for the identification of MARCH8 targets in human esophageal squamous cell carcinoma. J Proteomics 236, 104125.

Tada, T., Zhang, Y., Koyama, T., Tobiume, M., Tsunetsugu-Yokota, Y., Yamaoka, S., Fujita, H. and Tokunaga, K. (2015). MARCH8 inhibits HIV-1 infection by reducing virion incorporation of envelope glycoproteins. Nat Med 21, 1502–7.

Tobiume, M., Sato, Y., Katano, H., Nakajima, N., Tanaka, K., Noguchi, A., Inoue, S., Hasegawa, H., Iwasa, Y., Tanaka, J. et al. (2009). Rabies virus dissemination in neural tissues of autopsy cases due to rabies imported into Japan from the Philippines: Immunohistochemistry. Pathology International 59, 555–566.

Tze, L. E., Horikawa, K., Domaschenz, H., Howard, D. R., Roots, C. M., Rigby, R. J., Way, D. A., Ohmura-Hoshino, M., Ishido, S., Andoniou, C. E. et al. (2011). CD83 increases MHC II and CD86 on dendritic cells by opposing IL-10-driven MARCH1-mediated ubiquitination and degradation. J Exp Med 208, 149–65.

Umthong, S., Lynch, B., Timilsina, U., Waxman, B., Ivey, E. B. and Stavrou, S. (2021). Elucidating the Antiviral Mechanism of Different MARCH Factors. mBio 12.

van de Kooij, B., Verbrugge, I., de Vries, E., Gijsen, M., Montserrat, V., Maas, C., Neefjes, J. and Borst, J. (2013). Ubiquitination by the membrane-associated RING-CH-8 (MARCH-8) ligase controls steady-state cell surface expression of tumor necrosis factor-related apoptosis inducing ligand (TRAIL) receptor 1. J Biol Chem 288, 6617–28.

Yu, C., Li, S., Zhang, X., Khan, I., Ahmad, I., Zhou, Y., Li, S., Shi, J., Wang, Y. and Zheng, Y. H. (2020). MARCH8 Inhibits Ebola Virus Glycoprotein, Human Immunodeficiency Virus Type 1 Envelope Glycoprotein, and Avian Influenza Virus H5N1 Hemagglutinin Maturation. mBio 11.

Zhang, Y., Tada, T., Ozono, S., Kishigami, S., Fujita, H. and Tokunaga, K. (2020). MARCH8 inhibits viral infection by two different mechanisms. Elife 9.

Zhang, Y., Tada, T., Ozono, S., Yao, W., Tanaka, M., Yamaoka, S., Kishigami, S., Fujita, H. and Tokunaga, K. (2019). Membrane-associated RING-CH (MARCH) 1 and 2 are MARCH family members that inhibit HIV-1 infection. J Biol Chem 294, 3397–3405.

